# Evaluation of AlphaFold-Multimer prediction on multi-chain protein complexes

**DOI:** 10.1101/2022.12.08.519586

**Authors:** Wensi Zhu, Aditi Shenoy, Petras Kundrotas, Arne Elofsson

**Author notes:** The authors wish it to be known that the first two authors should be regarded as Joint First Authors.

## Abstract

**Motivation:** Despite near-experimental accuracy on single-chain predictions, there is still scope for improvement among multimeric predictions. Methods like AlphaFold-Multimer and FoldDock can accurately model dimers. However, how well these methods fare on larger complexes is still unclear. Further, evaluation methods of the quality of multimeric complexes are not well established.

**Results:** We analysed the performance of AlphaFold-Multimer on a homology-reduced dataset of homo- and heteromeric protein complexes. We highlight the differences between the pairwise and multi-interface evaluation of chains within a multimer. We describe why certain complexes perform well on one metric (e.g., TM-score) but poorly on another (e.g., DockQ). We propose a new score, Predicted DockQ version 2 (pDockQ2), to estimate the quality of each interface in a multimer. Finally, we modelled protein complexes (from CORUM) and identified two highly confident structures that do not have sequence homology to any existing structures.

**Availability:** All scripts, models, and data used to perform the analysis in this study are freely available at https://gitlab.com/ElofssonLab/afm-benchmark.

**Supplementary information:** Supplementary data are available at *Bioinformatics* online.

## 1. Introduction

Most biological processes and cellular functions depend on protein structures and their interactions. Understanding how proteins form three-dimensional structures can give critical insights into protein function. Therefore, the determination of the 3D structure of a protein from its primary amino acid sequence has been a fundamental problem in biology (Dill et al., 2008). With the recent advancement of AlphaFold (Jumper et al., 2021), obtaining a three-dimensional protein structure with near experimental accuracy is possible for most proteins by giving only the amino acid sequence as input. AlphaFold has been trained on protein chains and has shown remarkable performance in single-domain predictions. However, inside the densely packed cell environment, proteins are constantly near each other and perform functions by forming contacts with other biological macromolecules (Bergendahl & Marsh, 2017), often creating large biological complexes of multiple individual protein chains.

Predicting the structure of large molecular complexes has long been difficult unless a suitable template existed. Earlier methods used biophysical and biochemical interaction constraints (i.e. HADDOCK (Dominguez et al., 2003)) or pairwise docking predictions of components (i.e., Multi-LZerD (Esquivel-Rodríguez et al., 2012)) to model docking of protein-protein complexes with limited success. Shortly after the release of AlphaFold, it was recognised to help predict structures of protein assemblies if the input sequences are concatenated using flexible linkers or by modifying their residue numbers (Bryant et al., 2022; Mirdita et al., 2022). In FoldDock (Bryant et al., 2022) and ColabFold (Mirdita et al., 2022), the multiple sequence alignments (MSA) of the individual protein chains were combined by matching sequences based on the organisms (paired) and using block diagonalisation (block). The templates were disabled, and the combined alignments were submitted into the original AlphaFold pipeline. These studies showed that generating a “paired” alignment is crucial for protein-protein complex prediction. Simultaneously, AF2Complex (Gao et al., 2022) showed that using structural templates (without paired alignments) is often sufficient to predict structures of multimeric proteins. Methods like OmegaFold predict protein structures from a single primary amino acid sequence using protein language models without explicit MSAs (Wu et al., 2022). AlphaFold-Multimer, (Evans et al., 2022), an extension of AlphaFold for multimeric proteins, was specifically trained on multi-chains proteins.

In this study, we evaluated the performance of AlphaFold-Multimer predictions on a homology-reduced dataset independent from the AlphaFold-Multimer training set consisting of homomeric and heteromeric complexes with two to six chains. The model quality was evaluated against experimental structures using TM-score (Zhang & Skolnick, 2004) and DockQ (Basu & Wallner, 2016). While TM-score is more sensitive to global topology than local variations, DockQ assigns a higher weight to the accuracy of the predicted interface. The overall success rate ranges from ∼40-60% across all states, with a small decrease for larger heteromeric complexes. We also present a novel score, the second version of the predicted interface DockQ (pDockQ2), which estimates the quality of the interfaces in oligomers in the real-case scenario when the native (reference) structure is unknown and can be used to identify partially correct multimeric models.

## 2. Methods

### 2.1. Benchmark Dataset

Initially, the first biological unit of all structures with two to six chains (each with at least 30 residues) released after 30/04/2018 (the last date used in the training set of AlphaFold) was downloaded from the bio-units part of the Protein Data Bank (PDB). The structures were classified as homomers (all chains are 100% identical) or heteromers. Further, nine structures where at least one protein chain does not have contact with other proteins (this may occur after DNA/RNA removal from the PDB structure) were removed from the dataset, resulting in 23,222 proteins, with 13,136 dimers, 3,025 trimers, 4,257 tetramers, 942 pentamers, and 1,862 hexamers.

#### Similarity and homology reduction within each oligomeric state

We removed similar structures within each oligomeric state separately by aligning all-vs-all structures with MMalign (Mukherjee & Zhang, 2009) and subsequent clustering by the resulting MM-scores (TM-score (Zhang & Skolnick, 2005) calculated for all chains in the structure) utilising the Highly Connected Subgraphs (HCS) method (Hartuv & Shamir, 2000). We used an MM-score threshold of 0.6 for clustering, which roughly corresponds to the TM-score threshold for the individual proteins to have the same fold. To reduce the number of combinations, we only consider protein pairs where at least one protein is similar, using FoldSeek (van Kempen et al., 2022) with default settings. After clustering, the structure with the highest resolution was chosen as the representative structure. If two structures had the same resolution, the one with the minimum difference between the SEQRES sequence and ATOM section of the PDB record was selected, resulting in a dataset of 5,402 proteins, with 3,051 dimers, 548 trimers, 1,071 tetramers, 205 pentamers and 527 hexamers.

#### Homology reduction against the AlphaFold training dataset

MMseqs2 (Steinegger & Söding, 2017) removed homology between the benchmark dataset and the AlphaFold training dataset (PDB structures before 2018/04/30). A structure in our datasets was removed if at least one chain shared ≥ 30% sequence identity with any sequence in the AlphaFold training dataset. Finally, a manual examination of global stoichiometries for each oligomeric state was performed. Protein structures with conflicting stoichiometries were removed, resulting in 1,997 proteins, with 1,151 dimers, 224 trimers, 397 tetramers, 70 pentamers, and 155 hexamers. A subset of 837 complexes of various oligomeric states, for which Omegafold and ESMfold did not crash, was used to compare the performance with these methods.

### 2.2. CORUM Dataset

For further evaluation, we used the CORUM Version 3.0 Core Set (released in September 2018) (Giurgiu et al., 2019), containing 512 complexes with two to six chains. To identify complexes having no homology to any previous PDB structure, we ran MMseqs2 (Steinegger & Söding, 2017) between all chains of the selected CORUM complexes and PDB structures. If none of the chains in a CORUM complex has ≥ 30% sequence identity to any PDB chain, then this complex was kept for further modelling. This strict criterion reduced the number of CORUM complexes to 53, for which we ran through the AlphaFold-Multimer pipeline with its default settings. Out of those, 14 complexes did not produce MSAs for at least one of the chains, and in 10 cases, modelling failed due to out-of-memory or out-of-time errors, leaving 29 potentially novel complexes.

### 2.3. Model Generation

AlphaFold-Multimer (Evans et al., 2022) models were generated using AlphaFold v2.2.0 and the top-ranked model by AlphaFold was used for the analysis. The default parameters for multiple sequence alignment (MSA) generations and the number of recycles (3) were used. In 119 cases, the default pipeline did not produce all MSAs due to an out-of-memory error when run on 12 cores (Intel Xeon E5-2690v4) for 60 hours. These proteins were rerun using only one HHblits (Remmert et al., 2011) iteration and the reduced database small_bfd for 80 hours, leaving 18 proteins which did not generate alignments even with those reduced settings. AlphaFold-Multimer was run on a single NVIDIA DGX-A100 Core GPU for 72 hours. Still, the modelling of 60 complexes failed with the above settings due to an out-of-time error; these were ignored. The final dataset comprised 1928 protein complexes (1148 dimers, 220 trimers, 367 tetramers, 62 pentamers and 131 hexamers).

### 2.4. Evaluation

#### Scores to evaluate the quality of models against native PDB

We used two scores to evaluate the quality of the models - the DockQ score (Basu & Wallner, 2016) and the TM-score produced by the MM-align program (Mukherjee & Zhang, 2009) (henceforth referred to as MM-score). Both scores range between 0 and 1. According to the study (Basu & Wallner, 2016), if the DockQ score > 0.23, then the quality of the model is acceptable by the CAPRI criteria (Janin et al., 2003). MM-score was obtained using the default MM-align settings. It has been previously shown that when comparing structures of individual proteins, a TM-score of 0.5 roughly indicates the same fold. However, when considering a protein complex of two or more chains, it is possible to obtain scores higher than 0.5 if the larger chain(s) structure is correct. Still, the prediction of the quaternary structure can be wrong, i.e. it is necessary to use a higher cut-off to separate correct and incorrect multi-chain models.

### 2.5. DockQ for multimeric complexes

The DockQ score signifies the quality of an interface of a model compared with the native structure, with the larger protein acting as the receptor and the smaller protein as the ligand. In a multi-oligomeric complex, there are several ways to define an interface, e.g. (i) residues in contact between any pair of chains i and j with a complex, pairwise interface DockQ (DockQ_ij_) as illustrated in Fig 1A, or (ii) residues in contact between one chain i and all the other chains, interface DockQ (DockQ_i_) as illustrated in Fig 1B. Except for homomers with identical interfaces and dimers with only one interface, most cases have DockQ_i_ ≠ ∑_j_ DockQ_ij_. To ensure the order of the chains in the AlphaFold-Multimer model is the same as the native structure, we use the MMalign output alignment mapping chains between the native and modelled structures so that the chain names are consistent. For example, for a tetramer with chains A, B, C, and D, the DockQ_i_ was calculated using the following command line options:

**Fig 1.**
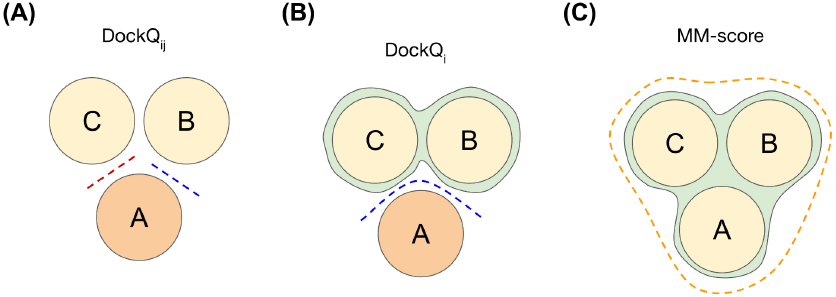
Schematic representation of two types of interface in an exemplary trimer (A) using pairwise interfaces (i.e. DockQ of chain A vs chain B and DockQ of chain A vs chain C) (B) using interface DockQ_i_ (i.e. DockQ when chain A is the ligand and chain B & chain C together form the receptor). Dashed lines in both panels represent interfaces with a non-zero number of contacts. (C) using overall structure as in MM-score.

~~~
python3 DockQ.py model.pdb native.pdb
   -model_chain1 A -native_chain1 A
   -model_chain2 B C D -native_chain2 B
   C D
~~~

while DockQ_ij_ was calculated for each pair of chains (A and B in the example) with the options

~~~
python3 DockQ.py model.pdb native.pdb
   -model_chain1 A -native_chain1 A
   -model_chain2 B -native_chain2 B
~~~

Note that the AlphaFold-Multimer models were generated for the full-length (PDB SEQRES section) sequences. Thus, to avoid the impact of residues, possibly not resolved experimentally, we calculated the DockQ score only for those residues present in both modelled and reference (PDB ATOM section) structures.

### 2.6 Predicting the quality of a model

AlphaFold-Multimer provides two intrinsic model accuracy estimates, pTM and ipTM. Both these scores estimate the average quality of the complex (or all interfaces of the complex), predicting the TM-score. However, in addition to estimating the quality of the entire predicted model, it is sometimes desirable to estimate each interface’s quality within a multi-chain complex. For dimers, (Bryant et al., 2022) proposed to use a predicted DockQ score (pDockQ) calculated from the number of contacts and the average quality of the interacting residues fitted to a sigmoid function:

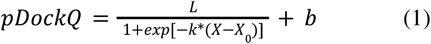

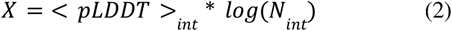

where <pLDDT>_int_ stands for the average of pLDDT (i.e. predicted lDDT score (Mariani et al., 2013) from AlphaFold-Multimer) over dimer interface residues, N_int_ is the number of interface contacts, and *X*_0_ and *b* are adjustable parameters. For multi-chain complexes, the pDockQ scores for chains i and j do not perform optimally (see Results). Therefore, we propose a novel variation of the predicted DockQ score, pDockQ2, using the relation

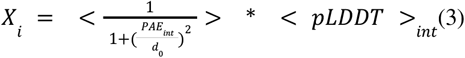

Here, PAE is the Predicted Aligned Error produced by AlphaFold-Multimer (Evans et al., 2022). The <PAE>_int_ is the PAE over all interfaces for chain *i*, first being scaled by an optimised parameter *d*_0_ = 10 Å. <pLDDT>_int_ is the average pLDDT of that combined interface residues. As in pDockQ, we fit a sigmoid curve (eq. 1) (by the scipy package) to the actual DockQ_i_ values, yielding the coefficients *L* = 1.31, *x*_0_ = 84.733, *k* = 0.075, and *b* = 0.005.

## 3. Results and Discussion

### 3.1. Interface quality in homomeric and heteromeric complexes using different measures

How to best evaluate the quality of a multimeric protein model is not well examined. Some metrics (e.g., MM-score) evaluate a complex in its entirety, while others provide evaluation per interface (DockQ_i_) or for each pair of proteins (DockQ_ij_). This study investigates correlations between these metrics types and the consistency within a complex. In Fig 2A, MM-score is compared to the maximum and minimum DockQ_ij_ scores for a complex and in Fig 2B, it is compared against DockQ_i_.

**Fig 2.**
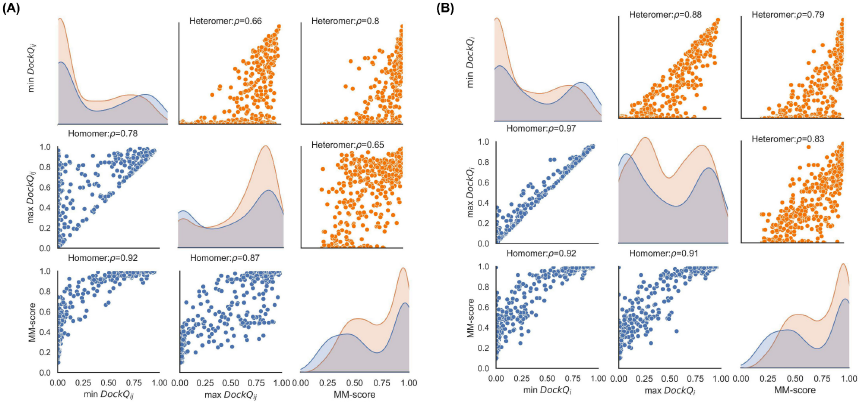
(A) Pairplot showing correlations between min DockQ_ij_, max DockQ_ij_ and MM-score for complexes > 2 chains (B) Pairplot showing correlations between min DockQ_i_, max DockQ_i_ and MM-score for complexes > 2 chains. Diagonal elements show density plots for corresponding quantities.

#### 3.1.1 Homomers vs Heteromers

For the heteromers, the spread of per-interface DockQ_*i*_ scores within a protein complex is larger than for homomers. Almost all homomeric complexes (80-90%) possess internal symmetry, repeating similar interfaces between subunits, leading to the uniform quality of the predicted interfaces and, thus, to a smaller variation in per-interface quality scores compared to heteromers, where each interface may be structurally unique. That explains the observation that for 97.5% of the complexes, all or none of the interfaces are predicted correctly in homomers (Fig 3A). For heteromers, one-fifth (20.5%) of the complexes have only a subset of the interfaces predicted correctly (Fig 3B).

**Fig 3.**
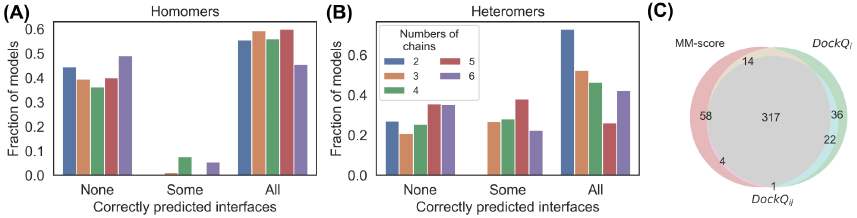
Fraction of models where none of the interfaces predicted correctly (NONE, i.e. no interface has DockQ_i_ ≥ 0.23), some of the interfaces predicted correctly (SOME, i.e. only some of the interfaces have DockQ_i_ ≥ 0.23) and where all of the interfaces predicted correctly (ALL, i.e. all the interfaces have DockQ_i_ ≥ 0.23) separately for each oligomeric state for homomers (panel A) and for heteromers (panel B). (C): Venn diagram for the number of successful docking models indicated by three different metrics(MM-score > 0.75, red circle, min DockQ_ij_ > 0.23, cyan circle, and min DockQ_i_ > 0.23, green circle).

#### 3.1.2 Variation of DockQ_i_ vs DockQ_ij_ within a complex

To perform per-complex comparisons, we aggregated per-interface scores (DockQ_ij_ and DockQ_i_) for an entire complex, using minimum or maximum scores for all interfaces. Variations in DockQ_ij_ values within a complex tend to be larger than in DockQ_i_. The average difference between min and max DockQ_i_ is 0.11, while the average difference between min and max DockQ_ij_ is 0.30 (Fig 2A vs 2B). Some examples of heteromers with partially correctly predicted interfaces and, consequently, big differences between min DockQ_ij_ and min DockQ_i_ scores are shown in Fig S 1.

A notable example is displayed in Fig S1A, where a significant difference in the scores is caused by a very low (only one in this case) number of native contacts, not predicted in the AlphaFold-Multimer model. In such cases, DockQ_ij_ for that interface is very low. In contrast, contributions of that interface to all DockQ_i_ scores are levelled out by correctly predicted interactions with other chains, yielding higher scores (for details, see caption to Fig S1).

Further, Fig 2B indicates that a significant fraction of the complexes have an MM-score indicating good overall modelling quality (>0.75) and low min DockQ_i_ score (<0.1), indicating that one protein chain is not correctly docked while the rest are correct. Out of these 58 complexes, only five are homomers, strengthening the earlier “all or none” observation for homomers. Fig 4A shows a homodimeric membrane protein (7STL) with a small alpha-helical swapped between the chains in the native structure. AlphaFold-Multimer predicts that those domains are associated with the corresponding main chain, i.e. not being domain-swapped. Fig 4B illustrates another homo-dimeric complex (6WKU) with low min DockQ_i_ and high MM-score for the AlphaFold-Multimer model. Here, the single protein chain contains a repeat of three domains. In the AlphaFold-Multimer model, the rotation between the chains is 120 degrees off, leading to a high MM-score as the structural alignment algorithm is performed without paying attention to the exact residue matching.

**Fig 4.**
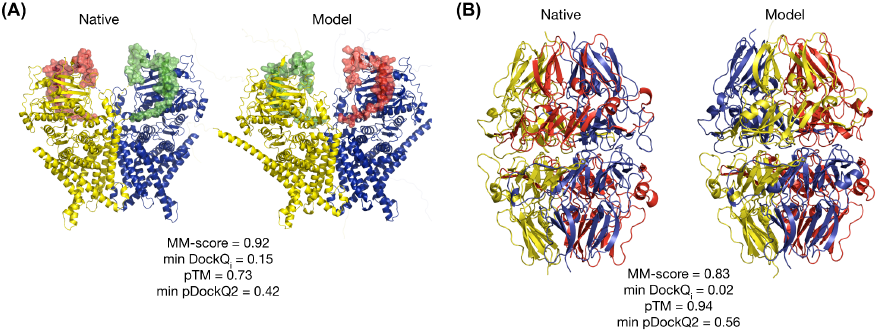
Different domain positions in the native (left panels) and model (right panels) structures. (A) Chains A and B of homodimer from PDB 7STL are shown as yellow and blue cartoons. Correspondingly, domains consisting of residues 868 - 914 are shown as semi-transparent green and red surfaces for chains A and B, respectively. (B) Chains A (lower structures) and B (upper structures) of a homodimer from PDB ID 6WKU. Each chain results from the fusion of three proteins with UniProt IDs Q01955 (residues 16 - 243, yellow cartoons), P53420 (residues 246 - 468, red cartoons), and P29400 (residues 471 - 695, blue cartoons). Disordered (flexible) parts of the model structures, which are missing in the native structures, but were modelled by AlphaFold-Multimer, are omitted for clarity.

Finally, we note a high overlap among the good models identified by the different methods using the following cutoffs, MM-score > 0.75, min DockQ_ij_ > 0.23, and min DockQ_i_ > 0.23, see Fig 3C. Overall, 70% of the good models are identified by all three measures, showing that for benchmarking, it is not crucial which measure is used. The min DockQ_i_ and max DockQ_i_ correlate significantly better (Spearman’s rank correlation coefficient, *r*=0.97) than the correlation between min DockQ_ij_ and max DockQ_ij_ (*r*=0.78). Moreover, our benchmarking study aimed to identify the models where all the chains and interfaces are completely correct (min DockQ_i_) instead of considering only the best possible interface (i.e. max DockQ_i_); hence all our subsequent analysis involves only the min DockQ_i_ for each complex.

### 3.2 Performance of AlphaFold-Multimer

Performance of AlphaFold-Multimer using MM-score and min DockQ_i_ for each oligomeric state are shown in Fig 5A and 5B. There is little difference between the homomers and heteromers, in general, using MM-score, whereas homomers have better accuracy if adopting min DockQ_i_ as the metric. The success rate for AlphaFold-Multimer for each oligomeric state, using min DockQ_i_ > 0.23, is reported in Fig 5C. Evaluations using per-interface measures can be found in Table S1-S2, and evaluations using per-complex measures can be found in Table S3-S5. The performance of AlphaFold-Multimer does not broadly vary across oligomeric states, meaning that, still, for complexes with six chains, there is an approximately 50% chance that the predicted protein complex is correct.

**Fig 5.**
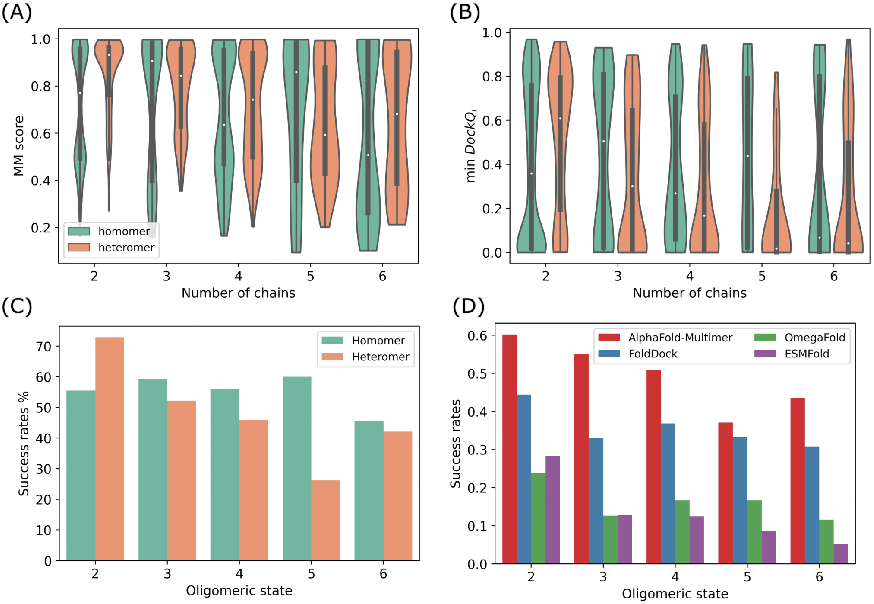
Performance comparison for different oligomeric states. (A),(B): AlphaFold-Multimer performance using(A) min DockQ_i_ and MM-score per complex between homomers and heteromers. The white dot represents the median, while the thick grey bar in the centre represents the interquartile range. The “violin shape” shows a kernel density estimation of the data. (C): Success rates (i.e. the fraction of acceptable models with min DockQ_i_ > 0.23) for AlphaFold-Multimer predictions. (D) Success rates on the common subset(n=837) of the benchmark dataset using AlphaFold-Multimer, FoldDock, OmegaFold, and ESMFold. The common subset comprises 592 dimers, 103 trimers, 110 tetramers, 6 pentamers and 26 hexamers.

Analysis of other multichain prediction methods, FoldDock (Bryant et al., 2022), Omegafold (Wu et al., 2022) and ESMFold (Lin et al., 2023; Wu et al., 2022) is conducted on a subset of our benchmark dataset 837 complexes of various oligomeric states, for which all four methods produced docking models. Fig 5D presents the success rate for each of the four prediction methods on that subset, and it shows that for dimers FoldDock makes around 40% correct models, while two language model-based methods OmegaFold and ESMFold, only make around 25% of such models (Table S6). FoldDock is worse than AlphaFold-Multimer for higher order multimers but still successfully docks 30-40% of the targets, while the corresponding numbers for OmegaFold and ESMFold are 10-15% and ∼10%.

In Fig S2, we examine the difference in quality for proteins included in the training set of AlphaFold-Multimer or not (deposited into PDB before 2018-04-30 (Evans et al., 2022)). The difference between homomers before and after the dataset is insignificant (P-value=0.49 obtained using Mann–Whitney T-test). However, heteromeric structures show a significant difference (P-value=1.98*10^−17^) with newer proteins being better than those used for training, i.e. there seems to be no indication that AlphaFold-Multimer is severely overtrained.

While distinguishing near-native (min DockQ_i_ > 0.23 and MM-score > 0.75) and incorrect docking models (min DockQ_i_ <=0.23 and MM-score < 0.75), we found that the number of effective sequences (N_eff_) in the paired alignment in wrong models is lower on average(Fig 6A). However, it is not an absolute causation. Although most complexes with min DockQ_i_ close to 0 have a N_eff_ score of 0, the inverse is false, i.e. the docking might fail even when the MSA is deep, Fig S3. Since we observe some heteromeric predictions with significant differences among chains, we check if the high variance among chain lengths within one complex might be a major factor for performance. Fig 6B shows no significant separation in chain length variance between good and bad models. Some failed cases have long disordered regions (i.e. residues in the SEQRES sequence missing from the ATOM record). However, some successful cases also have such long disordered regions, see Fig 6C. Symmetry does not appear to influence the quality of predicted models, see Fig 6D.

**Fig 6:**
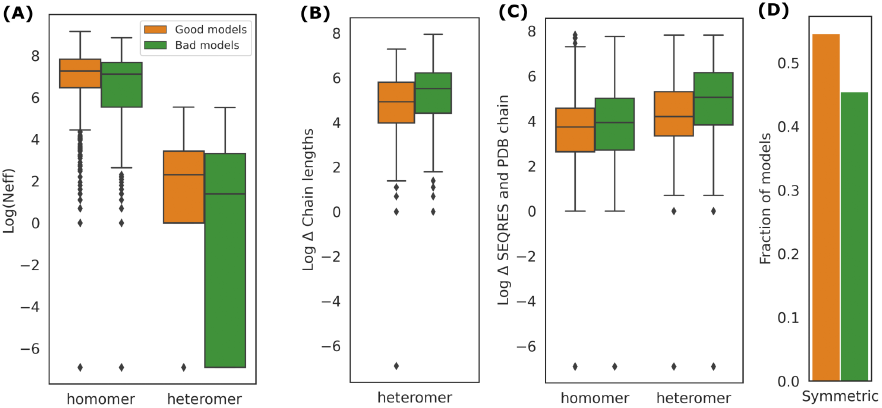
Comparison between models with (min DockQ_i_>0.23 and MM-score > 0.75, referred as “good models” and coloured in orange) and models with (min DockQ_i_ < 0.23 and MM-score < 0.75, referred as “bad models” and coloured in green) using (A) Log of the number of effective sequences (Neff) - small differences observed (p-value=9*10^−4^ for heteromers; p-value=1.4*10^−3^ for homomers). (B) Differences in length of chains within a heteromer - significant differences (p-value=1.75*10^−6^) (C) Differences between SEQRES (model) and PDB ATOM sequence (native) - significant differences observed (p-value=1.70*10^−30^ for heteromers; p-value=7.362*10^−6^ for homomers) (D) For symmetric protein complexes, the fraction of “good models” and “bad models”.

### 3.3 pDockQ2 - Improved estimate of DockQ for multi-chain predictions

In addition to per-structure quality estimation scores (pTM and ipTM), we believe there is a need to quantify the quality of each interface in a multi-chain complex prediction. Many models have a low min DockQ_i_ while a high pTM and ipTM, Fig S4. Next, we examined if pDockQ could be used. The results in Fig 7 indicate that pDockQ over-predicts DockQ_i_ for some complexes in this benchmark dataset. pDockQ was developed to predict DockQ on a dataset of heteromeric dimer models created with FoldDock. In Fig S5 and S6, we show that in datasets consisting of homomers or multimeric complexes and for models created with AlphaFold-Multimer, pDockQ sometimes gives high scores to incorrect models. More than 10% of the chains in all these sets have pDockQ > 0.5 and DockQ_i_ <0 .23. pDockQ does not utilise the predicted average errors (PAE) but only considers the size of the interface and the predicted quality (pLDDT) of residues in the interface. Therefore, it does not work if a method generates models with large, highly confident incorrect interfaces. To correctly classify such models as wrong, it is necessary to consider the PAE. Therefore, we developed pDockQ2 (see Methods), which considers PAEs between all chains. It is visually better correlated with DockQ_i_ for all subsets of models Fig 5, S5 and S6, except for heteromeric dimers using FoldDock, Fig S6B. A comparison between the two model confidence scores (pDockQ, pDockQ2) and DockQ_i_ shows that pDockQ2 also is better using FoldDock, Fig S7.

**Fig 7.**
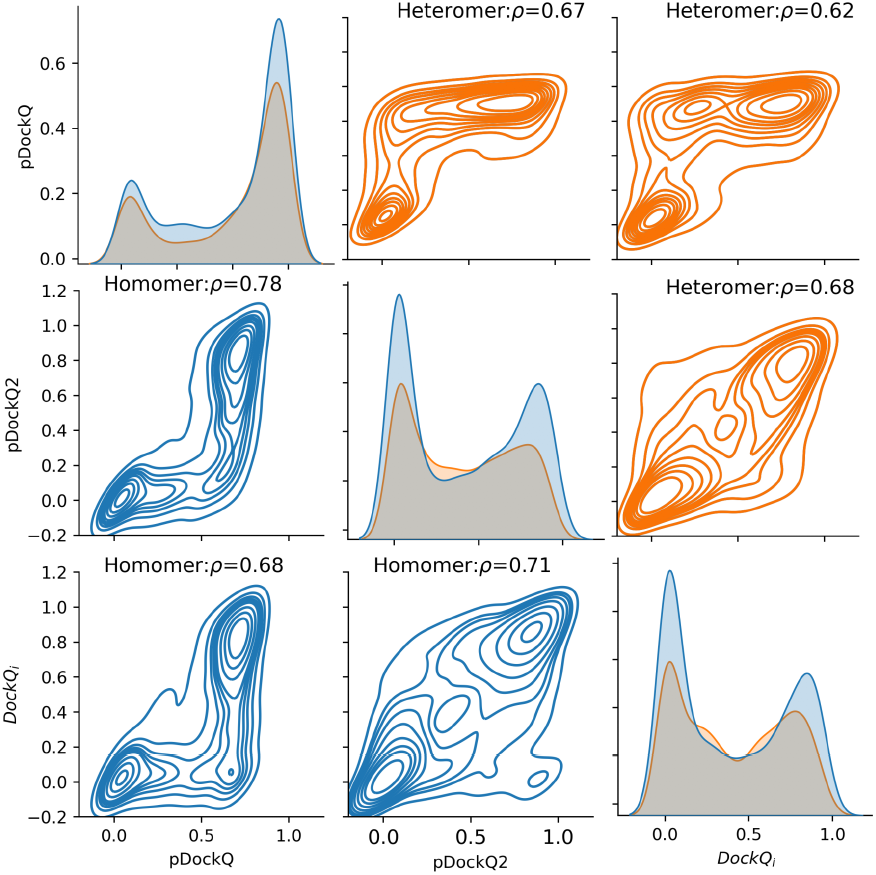
Pairplot showing the relationships between DockQ_i_ and predicted DockQ scores (pDockQ and pDockQ2) for interfaces of the complexes using AlphaFold-Multimer on the benchmark dataset. High confidence CORUM predictions

Although pDockQ2, in general, correlates well with DockQ_i_, for some models, the difference is large, i.e. some wrong models are predicted to be good or vice versa, and we took a further step on the possible reasons to explain these outliers. Using pDockQ2 < 0.1 and DockQ_i_ > 0.6 resulted in 55 interfaces. By comparing different metrics, we found that these models have a considerable length difference between the SEQRES sequences and the PDB sequences (p < 0.01 obtained using Mann–Whitney T-test) (see Fig S8). An example (PDB 6XWT) is shown in Fig S9A. The total SEQRES length of this hexamer is 2514 residues, whereas the PDB sequence length is only 352 residues. AlphaFold-Multimer successfully predicts the part of all chains which are aligned to the native structure well, and this produces min DockQ_i_ of 0.614. However, the overall estimation of the prediction gives low confidence, although pDockQ2 only utilises the residues in the interface. Further, 2 out of 6 interfaces have an average interface pLDDT below 50. Though the partial prediction might be correct, the confidence of the whole predicted model would be affected. Thus, the min pDockQ2 over all the interfaces for this prediction is 0.011.

We found that 72 predicted interfaces are the extreme cases with high pDockQ2 (>0.8) but low DockQ_i_ (<0.1), By manually checking those cases, the most prominent finding is that usually individual chains are predicted quite accurately, but the docking positions are wrong compared with the native structures. However, the high pDockQ2 values indicate that AlphaFold-Multimer is quite confident regarding the interface residues. Apart from this, 79% of the pTM scores for the complexes are above 0.8. Thus, other possible explanations might be needed. For example, the number of residues in contacts at the interface in the native structure of PDB 6JBD (Fig S5B) and PDB 5XLL (Fig S9C) is far fewer than those in the AlphaFold-Multimer prediction. We checked the original reference paper for the structure (Kita et al., 2020). We found that the dimer was cut out of a crystallisation structure of pantoate kinase, which has twofold homodimers. The AlphaFold-Multimer prediction gives the homodimer interface if one cuts the tetramer along the active binding site. In other words, the biological assembly from PDB might not represent the only biological assembly. In total, we encounter 65 such cases from 31 predicted models, which might be interesting to examine in detail, Table S7.

Now we asked if AlphaFold-Multimer can be used to predict truly novel structures by turning to the CORUM database. Out of the 29 potentially novel complexes (Table S8) that were predicted by AlphaFold-Multimer (see methods), 9 had pTM > 0.5 and min pDockQ2 > 0.23, see Fig S10. 2 highly confident complexes (pTM > 0.75 and pDockQ2 > 0.23) were obtained. Monomers of these complexes were then run with FoldSeek (van Kempen et al., 2022) against PDB. The PAC1-PAC2 complex (CORUM Complex ID: 3034) contains two chains, with chain A having a hit to 3GAA and chain B having a hit to 7LS6. However, the sequence identity is below 30% (see Fig S11A). The Metaxin complex (Mtx1, Mtx2) complex (CORUM Complex ID:3094) is also a dimer (Fig S11B) and is found in mice. Foldseek found a match to 6WUM with chain A and 6WUT to chain B. However, these hits (6WUM and 6WUT) are membrane proteins and part of the mitochondrial SAM complexes in Thermothelomyces thermophilus.

## 4. Conclusion

By preparing an independent homology-reduced dataset for benchmarking the performance of protein complex predictors, we have shown that taking the min DockQ_i_ over all interfaces is a useful way to evaluate the quality of the multimeric complex. Also, the performance of AlphaFold-Multimer (Evans et al., 2022) slightly decreases as the size of the complex increases, i.e. the number of chains in the complex increases, and consistent with the AlphaFold-Multimer study (Evans et al., 2022), homomeric complex prediction outperforms heteromeric complex prediction. By assessing the quality of the models with DockQ (Basu & Wallner, 2016) and MM-score (Mukherjee & Zhang, 2009), we show that for homomeric models, they are almost exclusively either completely correct or completely wrong, while for heteromeric complexes, there are cases where one or a few of the chains are incorrectly placed while the larger part of the complex is correct. We also provide a modified version of pDockQ, the pDockQ2 score for estimating the quality of an individual chain in a predicted multimer model. Lastly, we evaluate the chain-level predictions for highly confident structures using pDockQ2 obtained from CORUM using AlphaFold-Multimer. Here we present a structure of the Metaxin complex (Mtx1, Mtx2) complex having no detectable homology to any PDB structure.

## Author contributions

AS and WZ prepared the datasets and generated the models. All authors contributed to the analysis of results. AS wrote the initial draft of the manuscript. All authors contributed to the manuscript.

## Acknowledgements

We thank Gabriele Pozzati and Patrick Bryant for their valuable discussions.

