## Supplementary Information for "Evaluation of AlphaFold-Multimer prediction on multi-chain protein complexes"

### Supplementary Figures

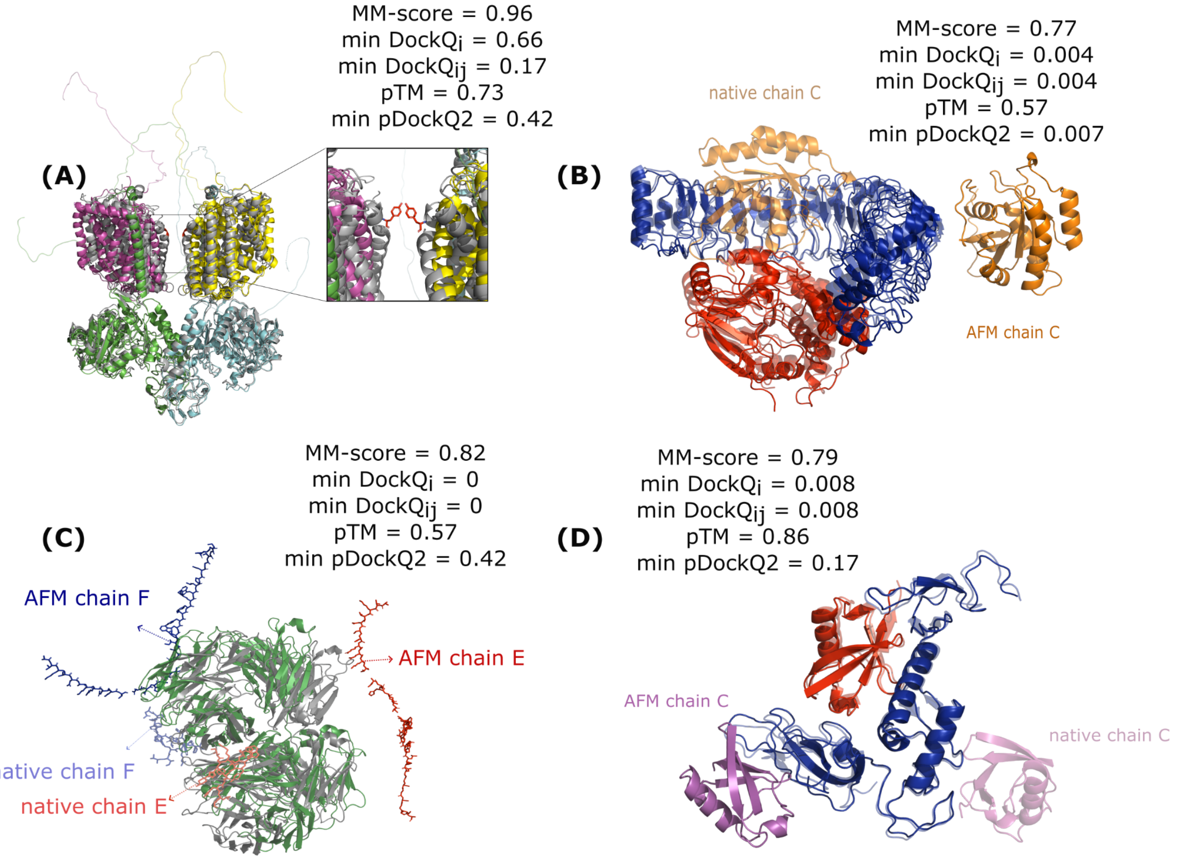


**Fig S1.** Examples of high variation in different evaluation scores. (a) PDB ID 6LI9 (tetramer) - shows the superimposition of modelled and native structure heteromeric amino acid transporter b0,+AT-rBAT complex bound with arginine (PDB ID: 6LI9). All four DockQ$_i$ give scores above 0.66. A single native contact (shown as sticks in the zoom-in window) between chains C and D was not predicted in the AlphaFold-Multimer model. Hence, the DockQ$_{CD}$ gives 0.17 for this interface despite the two chains looking perfectly aligned with the native structure. (b) PDB ID 7TXH (trimer) - shows the prediction of human MRas Q71R in complex with human Shoc2 LRR domain M173I and human PP1C (the native structure is blurred). The individual chains of this trimer are accurately predicted, and AlphaFold-Multimer only failed to put chain C in the correct position. For this predicted model, DockQ$_i$ from all three interfaces is below 0.11. However, since the interface between chains A and B is correct, the DockQ$_{AB}$ for this interface is 0.43. (c) 6CXG(hexamer). Chain A&B&C&D are in forest green in the AlphaFold-Multimer model, and the native structure is in grey. AlphaFold-Multimer puts chain E and chain F(glycopeptides) far away from the other four chains, so DockQ$_i$ involving these two chains are all 0. (d) 7US1 (trimer) Two out of three chains are correct, and the DockQ$_i$ for the prediction have minimum value 0.08 and maximum value 0.609. The MMscore is 0.790.

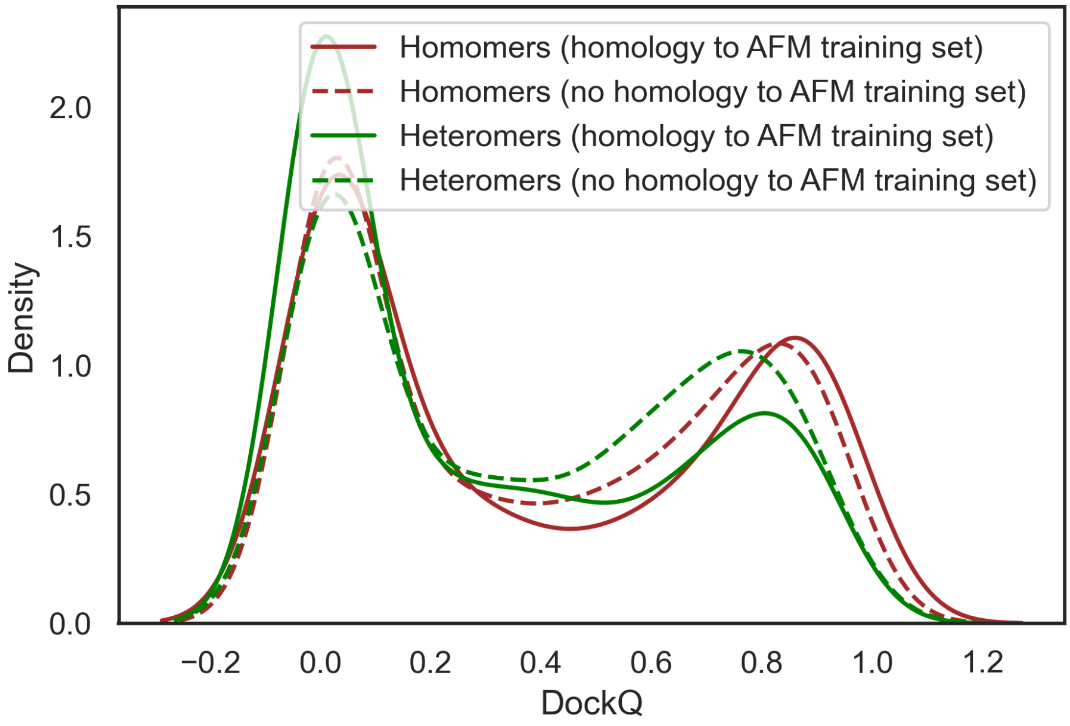


**Fig S2.** Comparison of DockQ distributions among homomeric and heteromeric complexes on the initial stage of our benchmark dataset, where the only difference is with or without the homology reduction between the AlphaFold-Multimer dataset. The homology reduction procedure is conducted by running HHblits with  E-value 10-3 against the UniClust30. If all the chains of a complex are homologous with an older complex (released before 2018-04-30), it is then removed from our dataset.

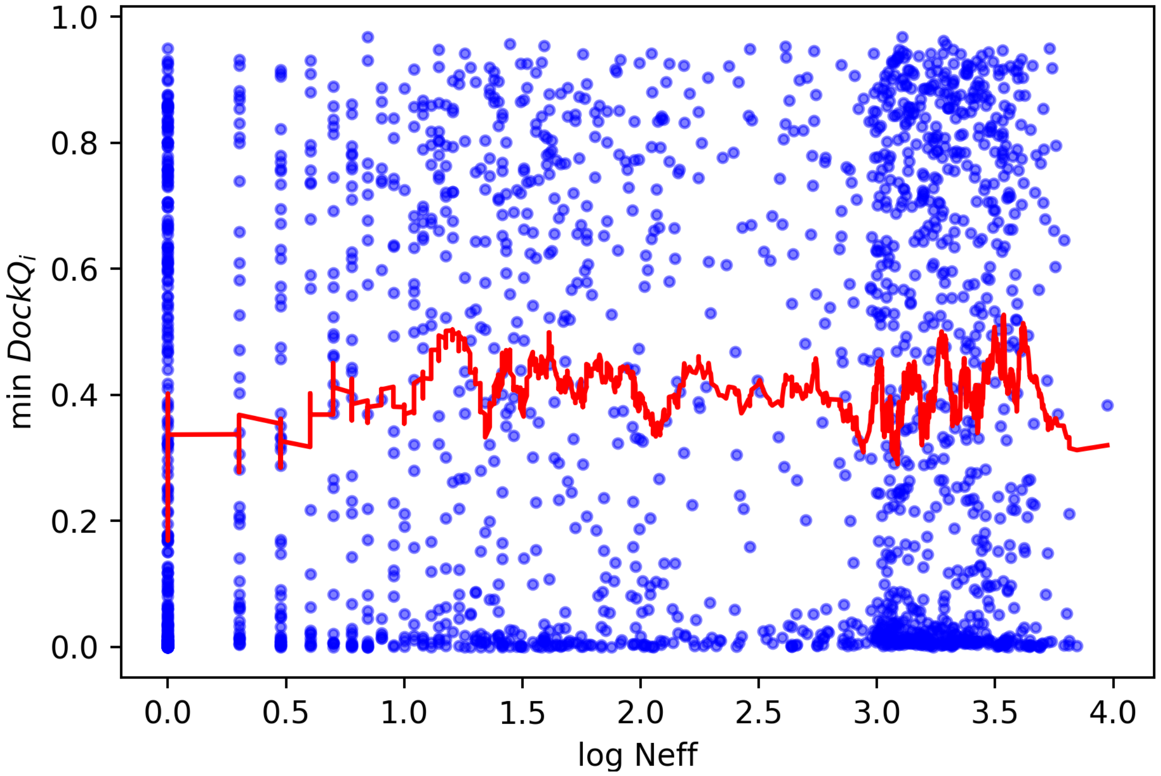


**Fig S3.** Scatterplot of min $DockQ_i$ and the logarithm of the number of effective sequences (Neff) of paired multiple sequence alignment for each protein complex. The red line shows the running average of min $DockQ_i$.

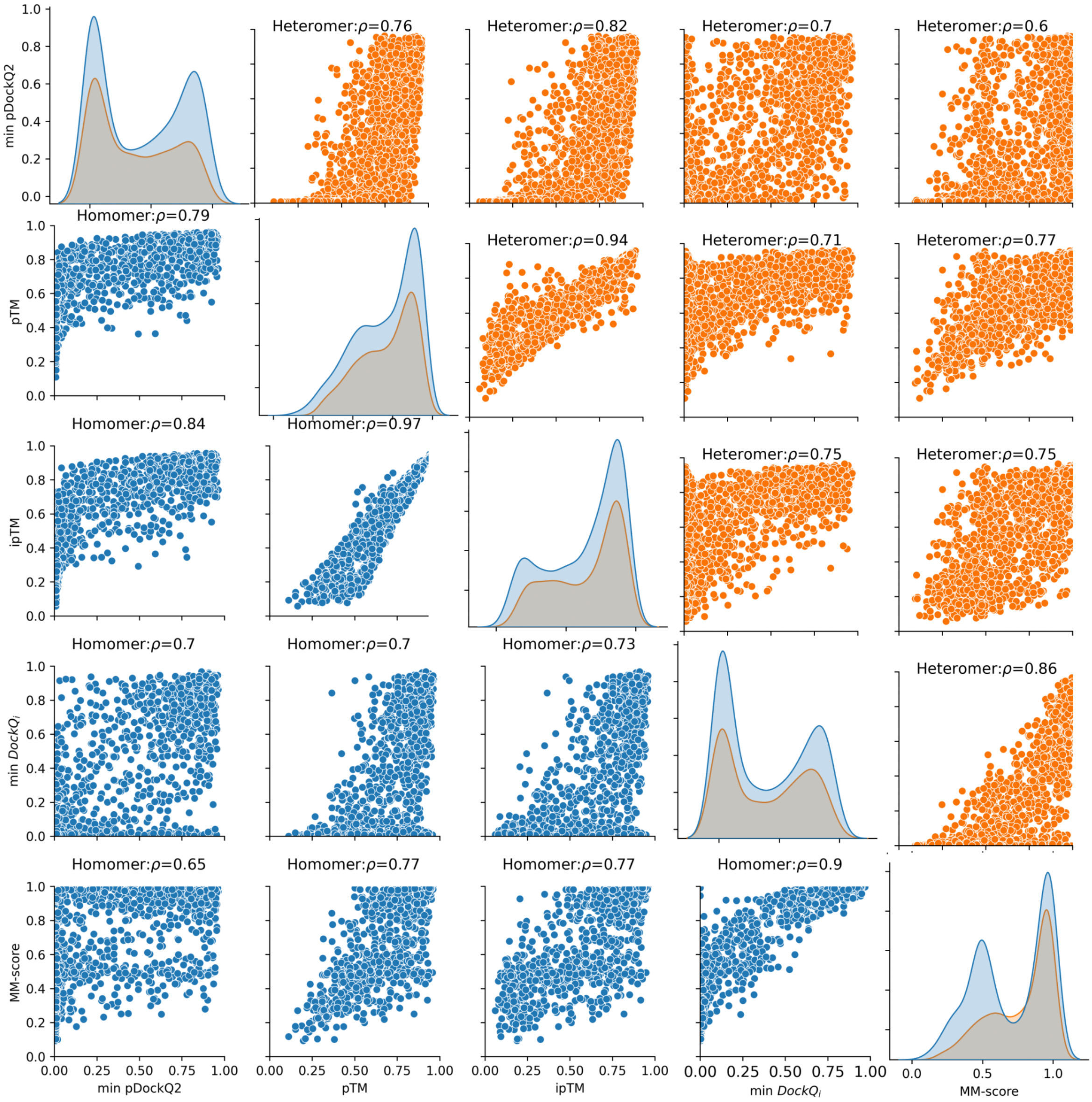


**Fig S4.** Pairplot showing the correlation between pDockQ2, quality measurements (min DockQ$_i$, MM-score) and confidence scores (pTM, ipTM).

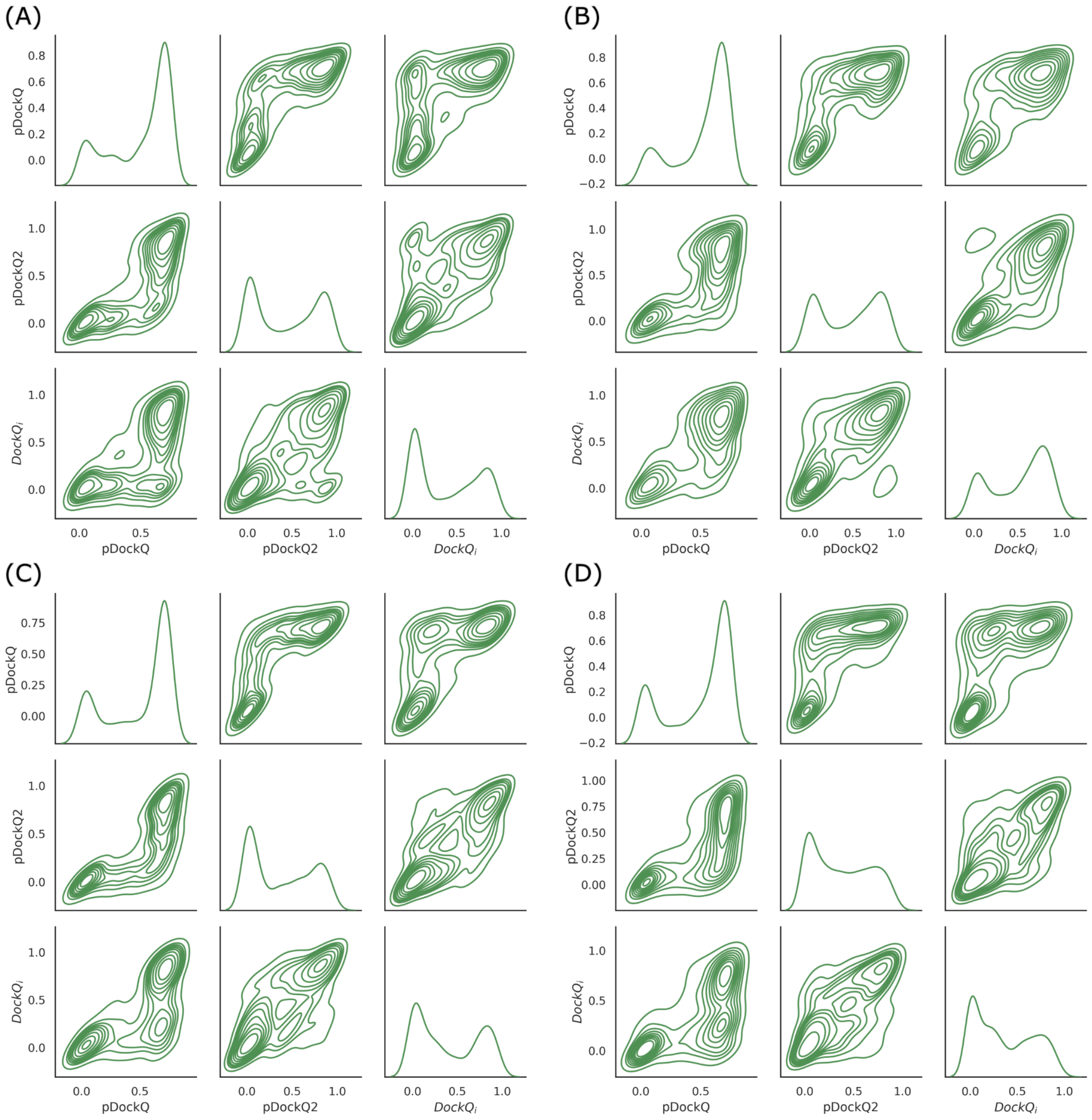


**Fig S5.** Pairplot representing the relationships between pDockQ, pDockQ2 and $DockQ_i$ for different oligomeric states using AlphaFold-Multimer on the common dataset(n=837). (A) homo-dimers. (B) hetero-dimers. (C) Homomeric protein complexes with three to six chains. (D) Heteromeric protein complexes with the three to six chains.

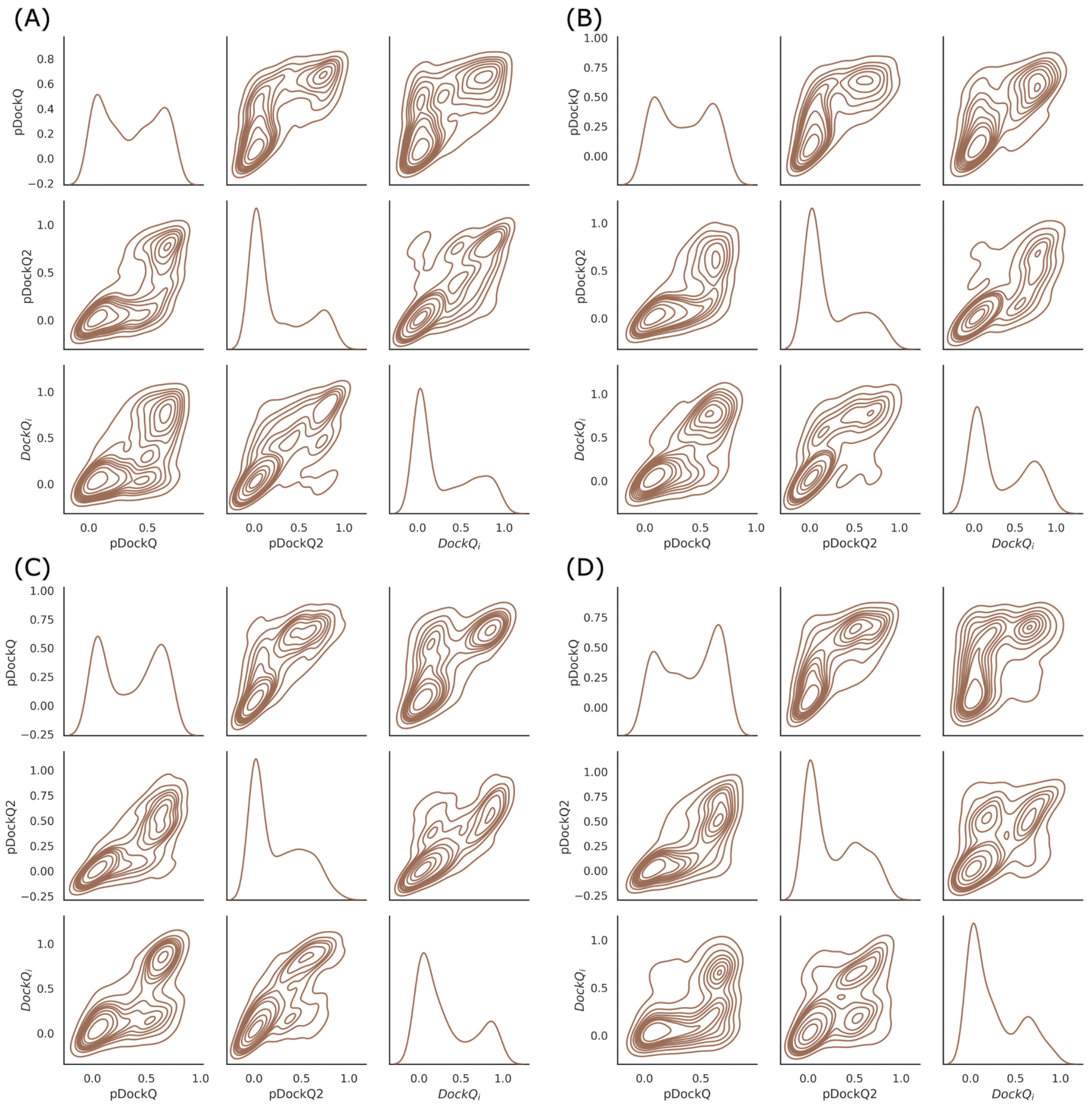


**Fig S6.** Pairplot representing the relationships between pDockQ, pDockQ2 and $DockQ_i$ for the different oligomeric states using FoldDock on the common dataset (n=837). (A) homo-dimers. (B) hetero-dimers. (C) Homomeric protein complexes with three to six chains. (D) Heteromeric protein complexes with three to six chains.

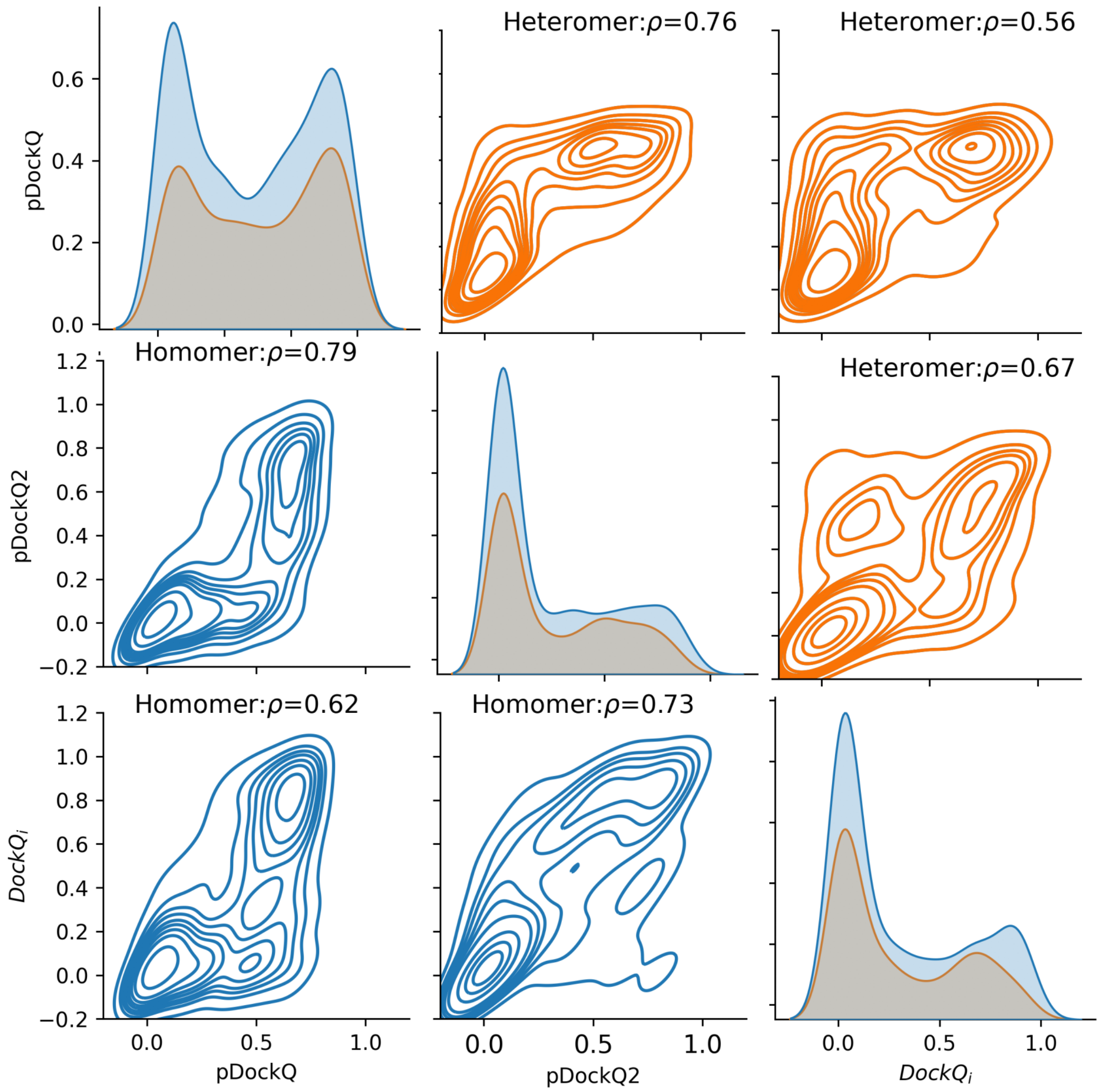


**Fig S7.** Pairplot showing the relationships between $DockQ_i$ and predicted DockQ scores (pDockQ and pDockQ2) for interfaces of the complexes using FoldDock on the benchmark dataset.

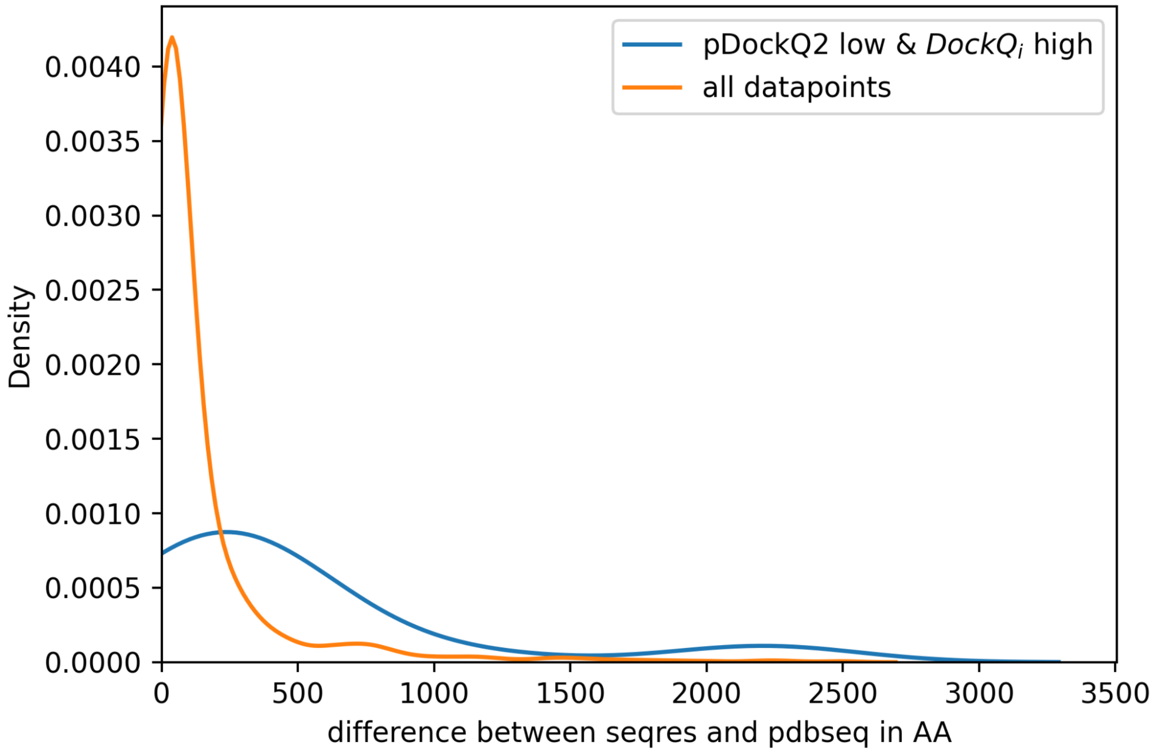


**Fig S8:** Distribution of difference in SEQRES and PDB chain length for complexes which fit pDockQ2 and those which have low pDockQ2 and high $DockQ_i$

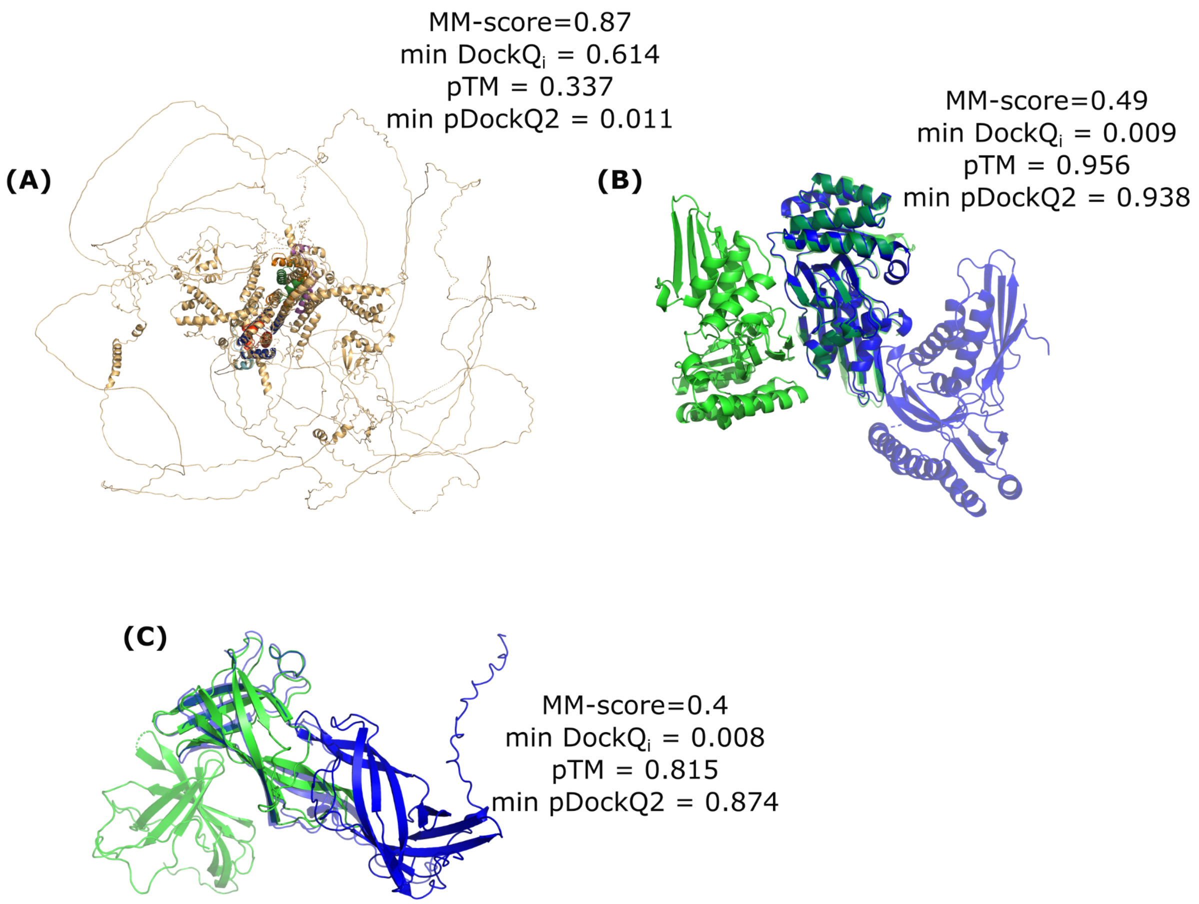


**Fig S9:** (A) PDB 6XWT, where the native structure is a subset of the model generated from the SEQRES sequence, and the model contains long disordered regions (B) PDB 6JBD - example with high difference in the number of interface contacts in native and model structure (C) PDB 5XLL where chains and native colour the model is coloured in blue and blurry.

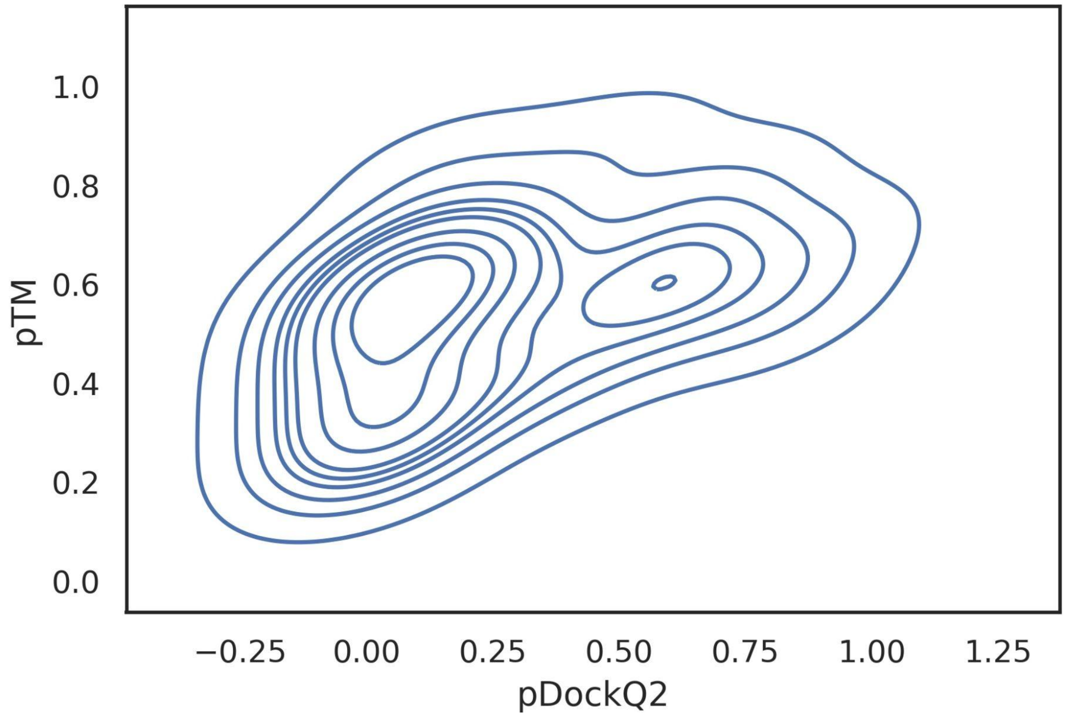


**Fig S10.** Kernel density plot between pTM (y-axis) and min pDockQ2 scores (x-axis) for the predicted CORUM complexes

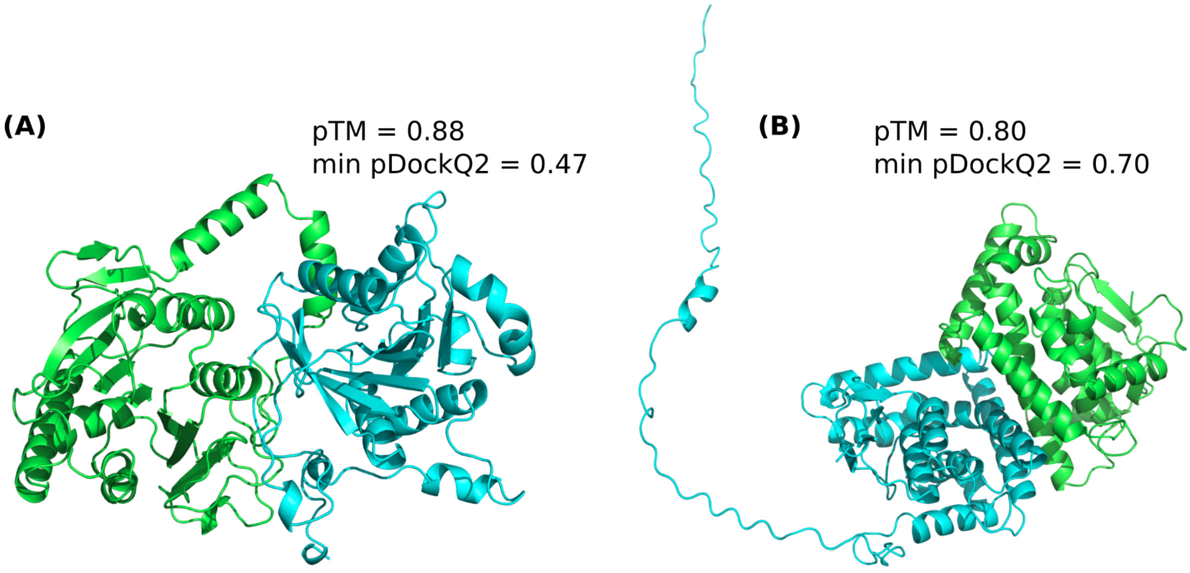


**Fig S11**. (A) Predicted structure of the PAC1-PAC2 complex (CORUM ID 3034) (B) Predicted structure for the Metaxin complex (Mtx1, Mtx2) (CORUM ID 3094)

### Supplementary Tables

**Table S1.** Success rates (i.e. the fraction of acceptable interfaces with $DockQ_i \geq 0.23$) for protein complex predictions by AlphaFold-Multimer

| | 2mer | 3mer | 4mer | 5mer | 6mer |
|---|---|---|---|---|---|
| Homomer | 55.5% (n=838) | 60.1% (n=86) | 60.4% (n=182) | 60.0% (n=20) | 47.0% (n=55) |
| Heteromer | 72.9% (n=310) | 65.9% (n=134) | 58.6% (n=185) | 42.4% (n=42) | 52.4% (n=76) |
| Overall | 60.2% (n=1148) | 63.7% (n=220) | 59.5% (n=367) | 48.1% (n=62) | 50.1% (n=131) |

**Table S2.** Success rates (i.e. the fraction of acceptable interfaces with $DockQ_{ij} \geq 0.23$) for protein complex predictions by AlphaFold-Multimer

| | 2mer | 3mer | 4mer | 5mer | 6mer |
|---|---|---|---|---|---|
| Homomer | 55.5% (n=838) | 57.7% (n=86) | 52.9% (n=182) | 45.7% (n=20) | 30.5% (n=55) |
| Heteromer | 72.9% (n=310) | 59.3% (n=134) | 46.5% (n=185) | 34.0% (n=42) | 36.3% (n=76) |
| Overall | 60.2% (n=1148) | 58.6% (n=220) | 49.7% (n=367) | 38.2% (n=62) | 33.6% (n=131) |

**Table S3.** Success rates (i.e. the fraction of acceptable models with min $DockQ_i \geq 0.23$) for protein complex predictions by AlphaFold-Multimer

| | 2mer | 3mer | 4mer | 5mer | 6mer |
|---|---|---|---|---|---|
| Homomer | 55.5% (n=838) | 59.3% (n=86) | 56.0% (n=182) | 60.0% (n=20) | 45.5% (n=55) |
| Heteromer | 72.9% (n=310) | 52.2% (n=134) | 45.9% (n=185) | 26.2% (n=42) | 42.1% (n=76) |
| Overall | 60.2% (n=1148) | 55% (n=220) | 50.9% (n=367) | 37.1% (n=62) | 43.5% (n=131) |

**Table S4.** Success rates (i.e. the fraction of acceptable models with max $DockQ_i \geq 0.23$) for protein complex predictions by AlphaFold-Multimer.

| | 2mer | 3mer | 4mer | 5mer | 6mer |
|---|---|---|---|---|---|
| Homomer | 55.5% (n=838) | 60.5% (n=86) | 63.7% (n=182) | 60.0% (n=20) | 50.9% (n=55) |
| Heteromer | 72.9% (n=310) | 79.1% (n=134) | 74.1% (n=185) | 64.3% (n=42) | 64.5% (n=76) |
| Overall | 60.2% (n=1148) | 71.8% (n=220) | 68.9% (n=367) | 62.9% (n=62) | 58.8% (n=131) |

**Table S5.** Success rates (i.e. the fraction of acceptable models with MMscore > 0.75) for protein complex predictions by AlphaFold-Multimer.

| | 2mer | 3mer | 4mer | 5mer | 6mer |
|---|---|---|---|---|---|
| Homomer | 50.6% (n=838) | 61.6% (n=86) | 44.5% (n=182) | 55% (n=20) | 40% (n=55) |
| Heteromer | 76.8% (n=310) | 64.9% (n=134) | 48.6% (n=185) | 38% (n=42) | 43.4% (n=76) |
| Overall | 57.7% (n=1148) | 63.6% (n=220) | 46.6% (n=367) | 43.5% (n=62) | 42% (n=131) |

**Table S6.** Success rates (i.e. the fraction of acceptable models with min $DockQ_i \geq 0.23$) for protein complex predictions by AlphaFold-Multimer, FoldDock, Haddock and OmegaFold for the common subset(n=837).

| Oligomeric state | AlphaFold-Multimer | FoldDock | OmegaFold | ESMFold |
|---|---|---|---|---|
| Dimer | 60.2% | 44.4% | 23.9% | 28.3% |
| Trimer | 55.0% | 33.0% | 12.6% | 12.7% |
| Tetramer | 51.0% | 36.8% | 16.7% | 12.4% |
| Pentamer | 37.1% | 33.3% | 16.7% | 8.7% |
| Hexamer | 43.5% | 30.8% | 11.5% | 5.1% |

**Table S7.** PDB IDs with high (>0.85) pDockQ2 (i.e. highly confident predictions by AlphaFold) and low (<=0.23) $DockQ_i$ scores. Possibly incorrect biological unit in PDB

| PDB ID | Number of chains | Class | MM-score | pTM | ipTM | Symmetry | min $DockQ_i$ | min pDockQ2 |
|---|---|---|---|---|---|---|---|---|
| 5xll | 2 | homomer | 0.4 | 0.815 | 0.794 | C2 | 0.008 | 0.874 |
| 5yek | 2 | homomer | 0.494 | 0.827 | 0.823 | C2 | 0.01 | 0.909 |
| 5ze7 | 2 | homomer | 0.489 | 0.937 | 0.925 | C1 | 0.006 | 0.922 |
| 6ea8 | 2 | homomer | 0.497 | 0.859 | 0.847 | C2 | 0.007 | 0.903 |
| 6eg7 | 2 | homomer | 0.494 | 0.899 | 0.905 | C2 | 0.013 | 0.887 |
| 6gf6 | 2 | homomer | 0.581 | 0.765 | 0.748 | C1 | 0.026 | 0.928 |
| 6jbd | 2 | homomer | 0.494 | 0.956 | 0.947 | C2 | 0.009 | 0.938 |
| 6k62 | 2 | homomer | 0.494 | 0.921 | 0.907 | C2 | 0.006 | 0.861 |
| 6kew | 2 | homomer | 0.491 | 0.896 | 0.876 | C1 | 0.002 | 0.911 |
| 6noy | 2 | homomer | 0.511 | 0.551 | 0.552 | C2 | 0.01 | 0.924 |
| 6sm4 | 2 | homomer | 0.498 | 0.763 | 0.718 | C2 | 0.016 | 0.919 |
| 6t4d | 2 | homomer | 0.507 | 0.861 | 0.856 | C2 | 0.006 | 0.93 |
| 6tgp | 2 | homomer | 0.508 | 0.86 | 0.84 | C1 | 0.03 | 0.926 |
| 6uxu | 2 | homomer | 0.498 | 0.936 | 0.915 | C2 | 0.006 | 0.911 |
| 6vd8 | 2 | homomer | 0.499 | 0.876 | 0.862 | C2 | 0.024 | 0.942 |
| 7cf7 | 2 | homomer | 0.428 | 0.915 | 0.897 | C2 | 0.025 | 0.936 |
| 7ebs | 2 | homomer | 0.475 | 0.929 | 0.915 | C2 | 0.008 | 0.864 |
| 7f9i | 2 | homomer | 0.523 | 0.726 | 0.703 | C1 | 0.042 | 0.945 |
| 7kpo | 2 | homomer | 0.279 | 0.673 | 0.712 | C2 | 0.01 | 0.885 |
| 7wsj | 2 | homomer | 0.48 | 0.784 | 0.741 | C2 | 0.025 | 0.903 |
| 5z5o | 2 | heteromer | 0.772 | 0.868 | 0.858 | C1 | 0.007 | 0.946 |
| 6dxo | 2 | heteromer | 0.526 | 0.709 | 0.76 | C1 | 0.223 | 0.922 |
| 6lki | 2 | heteromer | 0.614 | 0.898 | 0.878 | C1 | 0.02 | 0.898 |
| 6nep | 2 | heteromer | 0.643 | 0.894 | 0.869 | C1 | 0.014 | 0.87 |
| 6oqj | 2 | heteromer | 0.667 | 0.826 | 0.845 | C1 | 0.169 | 0.92 |
| 7b2i | 2 | heteromer | 0.507 | 0.919 | 0.939 | C1 | 0.006 | 0.875 |
| 7pku | 2 | heteromer | 0.603 | 0.781 | 0.851 | C1 | 0.22 | 0.852 |
| 6k6i | 3 | homomer | 0.332 | 0.926 | 0.915 | C3 | 0.01 | 0.9 |
| 6yaj | 4 | homomer | 0.594 | 0.8 | 0.759 | D2 | 0.188 | 0.931 |
| 7ddy | 4 | homomer | 0.743 | 0.897 | 0.891 | C1 | 0.009 | 0.954 |
| 7mq1 | 6 | homomer | 0.359 | 0.813 | 0.785 | D3 | 0.011 | 0.877 |

**Table S8.** Modelled CORUM IDs having no homology to existing structures

| CORUM ID | num_chains | Description | pTM | ipTM | min pDockQ2 |
|---|---|---|---|---|---|
| 12 | 3 | BLOC-2 (biogenesis of lysosome-related organelles complex 2) | 0.403 | 0.357 | 0.021 |
| 118 | 4 | GPI-GnT activity complex | 0.563 | 0.57 | 0.608 |
| 325 | 4 | - | 0.467 | 0.398 | 0.009 |
| 326 | 3 | - | 0.573 | 0.572 | 0.106 |
| 332 | 4 | - | 0.502 | 0.476 | 0.013 |
| 334 | 4 | - | 0.67 | 0.661 | 0.182 |
| 342 | 5 | - | 0.622 | 0.603 | 0.01 |
| 902 | 3 | - | 0.4 | 0.462 | 0.258 |
| 967 | 2 | - | 0.351 | 0.27 | 0.031 |
| 1057 | 2 | DIPA-MCRS1 complex | 0.281 | 0.094 | 0.009 |
| 1064 | 3 | IFP35-NMI complex | 0.608 | 0.527 | 0.028 |
| 1388 | 2 | - | 0.505 | 0.422 | 0.01 |
| 3020 | 2 | - | 0.22 | 0.143 | 0.008 |
| 3034 | 2 | PAC1-PAC2 complex | 0.879 | 0.886 | 0.47 |
| 3094 | 2 | Metaxin complex (Mtx1, Mtx2) complex | 0.798 | 0.86 | 0.702 |
| 3262 | 3 | SCAMP1-SCAMP2-SCAMP3 complex | 0.584 | 0.544 | 0.031 |
| 3525 | 4 | COG subcomplex (COG5, COG6, COG7, COG8) | 0.43 | 0.447 | 0.433 |
| 3532 | 2 | COG1-COG8 subcomplex | 0.685 | 0.705 | 0.622 |
| 3943 | 2 | - | 0.322 | 0.218 | 0.008 |
| 6407 | 2 | - | 0.667 | 0.656 | 0.115 |
| 6415 | 3 | pallidin-Cappuccino-BLOS1 complex | 0.585 | 0.619 | 0.425 |
| 6416 | 3 | dysbindin-Snapi | 0.536 | 0.684 | 0.834 |

| | | | | | |
|---|---|---|---|---|---|
| | | n-BLOS2 complex | | | |
| 6441 | 3 | KIAA0753-FOR20-OFD1 complex | 0.281 | 0.279 | 0.016 |
| 6458 | 2 | TIM23(sort) subcomplex (TIMM17B, TIMM23) | 0.683 | 0.744 | 0.878 |
| 6626 | 4 | AP5 adaptor complex | 0.615 | 0.576 | 0.189 |
| 6695 | 2 | MKKS-BBS12 complex | 0.829 | 0.898 | 0.108 |
| 6847 | 2 | DBIRD complex | 0.405 | 0.152 | 0.008 |
| 7411 | 5 | - | 0.609 | 0.648 | 0.521 |
